# Plasticity across levels: relating epigenomic, transcriptomic, and phenotypic responses to osmotic stress in a halotolerant microalga

**DOI:** 10.1101/2021.12.09.471842

**Authors:** Christelle Leung, Daphné Grulois, Luis-Miguel Chevin

**Affiliations:** CEFE, Université de Montpellier, CNRS, EPHE, IRD, Montpellier, France

**Keywords:** DNA methylation, gene expression, phenotypic plasticity, population dynamics, programmed-cell-death, RNA-sequencing, whole-genome bisulfite sequencing

## Abstract

Phenotypic plasticity, the ability of a given genotype to produce alternative phenotypes in response to its environment of development, is an important mechanism for coping with variable environments. While the mechanisms underlying phenotypic plasticity are diverse, their relative contributions need to be investigated quantitatively to better understand the evolvability of plasticity across biological levels. This requires relating plastic responses of the epigenome, transcriptome, and organismal phenotype, and how they vary with the genotype. Here we carried out this approach for responses to osmotic stress in *Dunaliella salina*, a green microalga that is a model organism for salinity tolerance. We compared two strains that show markedly different demographic responses to osmotic stress, and showed that these phenotypic responses involve strain- and environment-specific variation in gene expression levels, but a relative low - but significant - effect of strain × environment interaction. We also found an important genotype effect on the genome-wide methylation pattern, but little contribution from environmental conditions to the latter. However, we did detect a significant marginal effect of epigenetic variation on gene expression, beyond the influence of genetic differences on epigenetic state, and we showed that hypomethylated regions are correlated with higher gene expression. Our results indicate that epigenetic mechanisms are either not involved in the rapid plastic response to environmental change in this species, or involve only few changes in *trans* that are sufficient to trigger concerted changes in the expression of many genes, and phenotypic responses by multiple traits.

## 1. Introduction

Phenotypic plasticity, the ability of a given genotype to produce alternative phenotypes in response to its environment of development, is a prominent mechanism for coping with variable environments (Bradshaw 1965). Plasticity can evolve as an adaptation to spatially or temporally variable environments that are sufficiently predictable (Levins 1963; Via & Lande 1985; Gavrilets & Scheiner 1993; de Jong 1999; Tufto 2015). But plasticity may also be non-or maladaptive, leading to phenotypic change that has either no effect or detrimental effects, respectively, on fitness across environments (Ghalambor *et al*. 2007; Ghalambor *et al*. 2015). The adaptiveness of plasticity should ideally be assessed by comparing the fitnesses of plastic vs non-plastic genotypes across environments, but this is hardly feasible for most organisms and traits. An alternative is to focus on organisms for which it is clear that adaptive plasticity must be involved in their tolerance of a particularly broad and challenging range of environments.

An interesting example is provided by halotolerant organisms, such as the microalgae *Dunaliella salina*. This unicellular organism has the broadest range of salinity tolerance of all known eukaryotes, ranging in the laboratory from near freshwater to salt saturation (Ben-Amotz *et al*. 2009). This huge salinity range is afforded notably by outstanding physiological mechanisms of osmotic regulation (primarily through glycerol metabolism), great morphological flexibility, and unique specificities of its cell membrane (Ginzburg 1988; Pick 2002; Ben-Amotz *et al*. 2009). For these reasons, *Dunaliella* has been proposed as a model organism for investigating salinity tolerance in plants (Cowan *et al*. 1992). More generally, this also makes it a good model for studying adaptive phenotypic plasticity, as its phenotypic responses to salt have played a key ecological role in its evolutionary history, and were even shown recently to evolve in the laboratory in response to the predictability of environmental fluctuations, as predicted by theory (Leung *et al*. 2020).

Most existing predictions about the evolution of plasticity come from theoretical models that rely on simplifying assumptions about the mechanisms and inheritance of plasticity (Via & Lande 1985; de Jong 1990; Gavrilets & Scheiner 1993). These assumptions are made for mathematical convenience, but also out of lack of mechanistic understanding of plasticity. An important step towards more firmly relating these theoretical predictions to empirical measurements therefore involves deciphering the molecular mechanisms underlying phenotypic plasticity, its variation and inheritance. The molecular mechanisms contributing to phenotypic plasticity are diverse, but many of them rely on variation in gene expression with the environment (Beldade *et al*. 2011; Monteiro *et al*. 2015; Gibert *et al*. 2016). Such variation in gene expression may in turn be regulated by hormones and/or epigenetic processes, that is, a set of enzyme-mediated modifications resulting in the alteration of gene expression without any change in DNA sequences. Environmentally induced epigenetic variation has been proposed as a molecular mechanism underlying phenotypic plasticity (Angers *et al*. 2010; Bollati & Baccarelli 2010; Beldade *et al*. 2011). However, epigenetic variation can also result from other sources, most importantly genetic variation (Leung *et al*. 2016; Angers *et al*. 2020), and it is unclear to what extent plasticity in gene expression results from environmentally induced epigenetic variation.

To address these questions, we here investigated rapid responses to osmotic stress of two genetically divergent strains of the microalga *Dunaliella salina*, at three biological levels: the epigenome, the transcriptome, and macroscopic phenotypes. Not only does the outstanding ecology of this species make it ideal for studying plasticity, but it also has a reference genome since 2017 (Polle *et al*. 2017). This has recently led to work in comparative genomics aiming at identifying gene families involved in adaptation to salt (Polle *et al*. 2020), as well as investigations of plastic transcriptomic responses to salinity (Zhao *et al*. 2013; Fang *et al*. 2017; He *et al*. 2020), and specific epigenetic mechanisms such as small RNA (Lou *et al*. 2020). However, there has been no attempt to connect plastic responses to salinity at different levels (epigenome, transcriptome, and macroscopic phenotypes) across different genotypes, to decipher the underlying mechanisms of plasticity and their putative genetic variation. Our goal is thus twofold: (1) to further our understanding of the molecular mechanisms of plasticity in a model organism for salinity tolerance, including by shedding light on the (yet little investigated) contribution of DNA methylation; and (2) to more generally investigate how plastic and genetic variation are related at multiple levels of the organisms.

Our approach rests on comparing DNA methylation levels, gene expression levels, and demographic phenotypes, across environments and genotypes. Detecting a marginal effect of the environment on methylation patterns would confirm the environment as a source of epigenetic variation. Furthermore, considering epigenetic processes as an intermediate step between the genotype and the phenotype, we expected to detect a substantial contribution of the genotype to both epigenetic and phenotypic variation. And under the hypothesis that epigenetic processes play a role in fine-tuning gene expression, we expected to detect a correlation between the DNA methylation level and gene expression level. If epigenetic states and gene expression levels jointly change in response to environment, then this implies a role of epigenetics in gene-expression plasticity, which is thought to underlie the plasticity of most higher-level organismal traits (Beldade *et al*. 2011; Monteiro *et al*. 2015; Gibert *et al*. 2016). Finally, a significant genotype × environment interaction on gene expression and epigenetics would indicate an evolutionary potential for plasticity at these basal phenotypic levels of the organism.

## 2. Material and Method

### 2.1. Study strains and culture conditions

We compared two genetically related strains of *Dunaliella salina* (CCAP 19/12 and CCAP 19/15) originating from the Culture Collection of Algae and Protozoan (UK). For each of these strains, we obtained different replicates by using 10 different lines that were propagated independently in our laboratory for over two years (Leung *et al*. 2020; Rescan *et al*. 2020). Our standard growth conditions are 50 mL suspension flasks (CELLSTAR®; VWR 392-0016) containing artificial seawater with additional NaCl, complemented with 2% Guillard’s F/2 marine water enrichment solution (Sigma; G0154–500 mL), for a total 25 mL (including the inoculate), incubated at a constant temperature 24°C, and a 12:12 h light/dark cycle with a 200 μmol.m^-2^.s^-1^ light intensity. Target salinity was achieved by mixing the required volumes of hypo- ([NaCl] = 0 M) and hyper- ([NaCl] = 4.8 M) saline media, accounting for the salinity of the inoculate.

### 2.2. Population dynamics under osmotic stresses

To quantify salinity tolerance, we assessed the demographic responses of *Dunaliella salina* to three levels of salinity. To ensure similar physiological states and densities among all populations at the beginning of the population dynamics assays, we first performed an acclimation step during 10 days, by diluting all populations at 1:125 in fresh medium at intermediate salinity ([NaCl] = 2.4 M; note that this corresponds in absolute to a high salinity of *c*. 140g/L, but is considered as intermediate for our model species). We then inoculated *c*. 2 × 10^4^ cells.mL^-1^ of each populations into low (0.8 M), intermediate (2.4 M) or high (4.0 M) salinity, and tracked population density for the next 10 days under our standard growth conditions. We also followed the population dynamics of six randomly chosen populations of CCAP 19/15 strain starting at a lower density of 5 × 10^3^ cells.ml^-1^, to assess the effect of initial density on population growth rate in hyper-osmotic condition ([NaCl] = 4.0 M).

To measure population growth rates under the different salt concentrations, we assessed population densities by passing a subsample of 150 μl of each populations through a Guava® EasyCyte™ HT flow cytometer (Luminex Corporation, TX, USA), following the protocol described in Leung *et al*. (2020). Discrimination between alive and dead algae was possible thanks to chlorophyll auto-fluorescence detected through a cytogram of emissions at Red-B (695/50 nm) and Yellow-B (583/26 nm) band pass filters (Papageorgiou 2004), and the particle size was assessed through the Forward Scatter (FSC) and Side Scatter (SSC) parameters (Adan *et al*. 2017). To estimate population dynamics, cell counts were performed for alive algae at 11 time points: end of the acclimation step, 4h after the transfer to fresh media, and once per day for the following nine days.

### 2.3. Sample preparation, sequencing, and bioinformatic preprocessing

To investigate the molecular mechanisms involved in osmotic stress responses, we performed whole-transcriptome shotgun sequencing (RNA-seq) for the comparison of gene expression levels, and whole-genome bisulphite sequencing (WGB-seq) for the comparison of DNA methylation variation among the two strains (CCAP 19/12 and CCAP 19/15), in two environmental conditions (hypo- and hyper-osmotic). At the end of an acclimation step as described above, we transferred two biological replicates per strain to low ([NaCl] = 0.8 M) and high ([NaCl] = 4.0 M) salinities, in a greater volume (250 ml) than for the demographic assays, and at a density of *c*. 1 × 10^5^ cells.mL^-1^ so as to ensure enough material for high-throughput sequencing. After 24h following the salinity changes, the microalgae cells were harvested by centrifugation at 5,000 rpm for 15 minutes at room temperature, and cell pellets were stored at - 80°C until acid nucleic extraction.

Total RNA extraction and purification of the eight samples (2 strains × 2 salinities × 2 replicates) was carried out using Nucleozol® following Macherey Nagel’s protocol, and whole genomic DNA was isolated according to the phenol-chloroform purification and ethanol precipitation method of Sambrook *et al*. (1989). Library construction (TruSeq RNA Library Preparation kit for RNA-seq and Swift Bioscience Accel-NGS Methyl-Seq DNA library Kit for WGB-seq) and high-throughput sequencing steps (Paired-End (PE) 2□×□150□bp, Illumina® HiSeq®) were performed by Genewiz (Leipzig, Germany). We then performed all the bioinformatic preprocessing analyses with publicly available software implemented in the European UseGalaxy server (Afgan *et al*. 2018).

### Gene expression analyses

The RNA-seq raw reads were checked for quality using *FastQC* version 0.72 (Andrews 2010) and subjected to adapter trimming and quality filtering using *Trim Galore!* version 0.4.3.1 (Krueger 2015). Additional 12 bp and 3 bp were also removed at the 5’ and 3’ extremity, respectively, to avoid bias not directly related to adapter sequences or basecall quality according to *FastQC* outputs, and only reads with a minimum length of 50 bp were retained. We aligned the trimmed reads on the *D. salina* CCAP 19/18 reference nuclear (Dunsal1 v. 2, GenBank accession: GCA_002284615.2), chloroplastic (GenBank accession: GQ250046) and mitochondrial (GenBank accession: GQ250045) genomes, using *HISAT2* version 2.1.0 (Kim *et al*. 2015a) with default parameters for PE reads and spliced alignment option. We used *Stringtie* version 2.1.1 (Pertea *et al*. 2015) to predict transcript structures of each library based on the aligned reads, and performed *de novo* transcriptome assembly using the *Stringtie merge* tool, thus generating a unified and non-redundant set of transcripts across the different RNA-seq samples. We finally quantified the number of reads per transcript with *FeatureCounts* version 2.0.1 (Liao *et al*. 2014) using the alignment files from *HISAT2* and the transcript annotation file from *Stringtie*.

### Genetic variant calling and genetic diversity analysis

We assessed the genetic differences between strains based on the sequences from the transcriptomic data. We first calculated the genomic range from which variant calling was performed as the number of bases in all exons, after merging any overlapping exons from different transcripts. BAM files from previous read alignment analyses with *HISAT2* were submitted to *freebayes* (Garrison & Marth 2012) for variant identification, with the following parameters: joint variant calling of the eight samples simultaneously, minimum alignment quality of three, ploidy set to one, and samples assumed to result from pooled sequencing. After left-alignment and normalization of indels using *bcftools* (Li *et al*. 2009), we filtered the obtained *freebayes* multi-sample VCF file for strand bias (SPR and SAP > 20), placement bias (EPP > 20), variant quality (QUAL > 30), and depth of coverage (DP > 20). To assess genetic diversity within and differentiation between strains, we calculated the gene diversity for each library (H_E_), mean gene diversity for each strain (H_S_), total gene diversity (H_T_), and genetic differentiations (G_ST_) among samples within each strain, and also between the two strains for all variants. We then represented the Euclidean genetic distances among samples with a dendrogram computed with *ape R* package (Paradis & Schliep 2019).

Finally, for each locus that diverged between the strains (i.e., with alleles nearly fixed within strain but different between strains, H_S_ ≤ 0.1 and G_ST_ ≥ 0.9), we assessed the synonymous vs non-synonymous status of the substitution relative to the reference genome by using *SnpEff* version 4.3 (Cingolani *et al*. 2012). Sites where one of the focal strains had the reference allele while the other strain exhibited a synonymous mutation relative to the reference genome represented synonymous substitutions between our focal strains. As we detected only a single locus displaying a non-synonymous mutation in both strains relative to the references, we did not consider it in the synonymous vs non-synonymous mutation comparison between the strains.

### Methylation calling

The WGB-seq raw reads were also checked for quality using *FastQC*. Adapter and low-quality sequences were then trimmed using *Trim Galore!* Version 0.4.3.1. As specified by the Accel-NGS Methyl-seq Kit manual, additional 15 bp and 5 bp were trimmed at the 5’ and 3’ extremity, respectively, to remove the tail added during library preparation, thus avoiding non-quality-related bias. Mapping was performed on the same references genomes as in RNA-seq analyses, using *Bismark Mapper* version 0.22.1 (Krueger & Andrews 2011). Only uniquely mapping reads were retained, using *Bismark Deduplicate* tool. We then extracted the methylation status from the resulting alignment files using *MethylDackel* (Galaxy Version 0.3.0.1), where only cytosines covered by a minimum of 10 reads in each library were considered, and with the option of excluding likely variant sites (i.e. minimum depth for variant avoidance of 10×, and maximum tolerated variant fraction of 0.95). Cytosine methylation levels were determined for each CpG, CHG and CHH context. The high bisulfite conversion rate (> 99%) was assessed by Genewiz by spiking in unmethylated lambda DNA in three randomly chosen libraries, and it was also confirmed by our analyses by estimating the number of methylated cytosine calls in the organelle genomes.

### Statistical analyses

#### Population dynamics

To investigate how the population dynamics of our strains varied in response to salinity, we performed General Linear Models (GLMs) using cells count data as responses variables, with a negative binomial distribution (and log-link function). We performed GLMs on population size using strain, salinity (treated as categorical variables), day (as continuous variable), and their interactions as fixed effect. In such models with logarithmic link functions, any effect of time (here day) translates into a rate of exponential growth (or decline if negative) per unit time (here per day), and interactions of time with other factors estimate effects on population growth, which are our main interest. We performed such models on initial growth rate (Day 0 to Day 1: GLM #1) and growth rate in the exponential phase (Day 1 to Day 4: GLM #2). We also investigated whether difference in growth rates could be explained by a release of density dependent competition. To do so, we tested the effect of initial density on population growth rate of one of the strains in hyper-osmotic condition, using populations starting at 20,000 or 5,000 cells.mL^-1^ (Day 1 to Day 4 at [NaCl] = 4.0 M: GLM #3 in Table S2). Statistical analyses were conducted using the statistical environment R version 4.0.3 (R Core Team 2020) with the *MASS* package (Venables & Ripley 2002) for the GLMs.

### Differential gene expression and DNA methylation analyses

The differential expression analyses were performed with the Bioconductor’s package *DESeq2* version 1.30.1 (Love *et al*. 2014). We first normalized the count matrix using *DESeq2* regularized log transformed (*rlog*). We applied a redundancy analyses (RDA (Borcard *et al*. 1992)), computed with the function *rda()* available in the *vegan* R package (Oksanen *et al*. 2020), to quantify the proportions of the total gene expression variation that are significantly explained by strain, salinity or the strain × salinity interaction, with population identity as covariates to account for paired samples between salinities. We also identified differentially expressed transcripts among the same three factors, by building a general linear model as implemented in *DESeq2* and using Wald significance tests (Love *et al*. 2014). Transcripts with FDR < 0.05 (*P*-values after Benjamini-Hochberg (BH) adjustment) and |log_2_FC| > 1 were considered as differentially expressed. Significance of the strain × salinity interaction term was assessed using likelihood ratio tests (LRT) comparing models with and without the interaction term (Love *et al*. 2014).

Similarly, we performed a RDA to quantify the proportions of the total variation in DNA methylation level that are significantly explained by strain, salinity or strain × salinity interaction. We then used Bioconductor’s *methylKit* package (Akalin *et al*. 2012) to identify differentially methylated regions (DMRs), corresponding to non-overlapping 100 bp windows between strains, or between salinities for each strain. The significance of calculated differences was determined using Fisher’s exact tests. We used the Benjamini-Hochberg (BH) adjustment of *P*-values (FDR < 0.05) and methylation difference cut-offs of 20%.

For both RNA-seq and WGB-seq data, we performed Principal Component analyses (PCA) to represent the total variation among samples along its major axes. We also illustrated the expression level of DE transcripts and methylation levels of identified DMRs of all samples with heat-maps.

### Correlation between DNA methylation and gene expression

Since gene expression is a crucial step in the mapping from genotypes to phenotypes, and is thought to be a key mechanism underlying phenotypic plasticity, we wished to quantify to what extent gene expression levels can be predicted by key covariates in our dataset. We first evaluated the different sources of variation in gene expression by achieving a global partitioning of variation, assessing the influence of i) the genotype, ii) the environment, and iii) epigenetic variation on the total gene expression. We performed a RDA using the *rlog* transformed transcript count table as the response variable, and strain identity, salinity and epigenetic variation as explanatory variables. To account for paired samples among salinities, partial RDA was performed by removing the effect of population identity prior to assessing the effect of the environment or epigenetic variation on gene expression variation. Prior to RDA, we performed a PCA on DNA methylation levels and used the principal component (PCs) factors explaining at least 10% of the variation as a multivariate summary of epigenetic variation. We quantified the contributions to the total gene expression variation using the adjusted *R*^*2*^, and tested the significance of each *R*^*2*^ by ANOVA-like permutation tests using 999 randomization of the data (Legendre & Legendre 1998).

To further quantify the role of DNA methylation in gene expression, we correlated differences in local DNA methylation to fold-changes in expression of the corresponding transcript. In the absence of well-annotated *D. salina* genome, we associated each cytosine to a given transcript based on its distance to the nearest transcription start site (TSS). Using the *genomation* package (Akalin et al. 2015), we first calculated TSS coordinates using the gene structure file from the *de novo* transcriptome assembly of the RNA-seq preprocessing analyses. We then got the distance to nearest TSS and associated transcript identity for each cytosine. All cytosines associated to the same transcript were merged into a common gene-associated methylation region. We identified methylation differences between strains, and between salinities for each strain, using the same criterion for these gene-associated methylation regions as for DMRs from sliding windows above (i.e. FDR < 0.05 and |ΔmCG| > 20%). We finally compared the mean expression fold change associated to hypo- (< -20%) or hyper- (> 20%) methylated DMRs for a given comparison. We restricted our correlation analysis to genomic regions that were both detected as significantly methylated between strains or salinities, and associated to transcripts that also showed significant differential expression between strains or salinities.

## 3. Results

### 3.1. Genetic differences between strains

RNA-sequencing generated 1.92 × 10^8^ 150bp paired-end raw reads from eight samples (Table S1). The genomic range from which variant calling were processed represented a total of 4.043 × 10^7^ nucleotides. Across these loci, we identified a total of 6,201 (0.015%) variants among all samples. Among these polymorphic loci, we detected 5,500 loci displaying synonymous substitutions between the strains. This represents a moderate synonymous divergence of *c*. 10^−4^ per base pair between the two strains, consistent with previous observations from ITS sequences that led to placing these strains closeby in the *Dunaliella* phylogeny (Assunção *et al*. 2012; Emami *et al*. 2015). Where this has been investigated (in animals), such a level of synonymous divergence is consistent with within-rather than between-species variation (Roux *et al*. 2016), even though this notion becomes less clear for highly clonal micro-organisms.

Despite this low divergence, genetic variation was highly structured between the strains. Across the 6,201 variants, 6,170 (99.50%) displayed only two alleles across all the populations, and we measured a very low mean genetic diversity for these variants within each sample of a given strain (H_S_ = 0.03, *sd* = 0.097 and 0.039, *sd* = 0.102 for CCAP 19/12 and 19/15, respectively, Fig. S1-A and B), but a very high total genetic diversity across all samples (H_T_ = 0.439, *sd* = 0.313, Fig. S1-C), indicating highly structured genetic variation across strains (Fig. 1). Specifically, 5,589 (89.97%) variants displayed a fixed allele within a given strain, but distinct alleles between the strains (H_S_ ≤ 0.1 and G_ST_ ≥ 0.9). While we confirmed the genetic differences between the two strains, we also observed that samples of the same strain are quite genetically identical (Fig. 1 and Fig. S1-D and E). As a consequence, in subsequent analyses we used strain identity as a qualitative explanatory factor for genetic effects, instead of quantitative values based on genetic distances among samples.

**Fig. 1.**
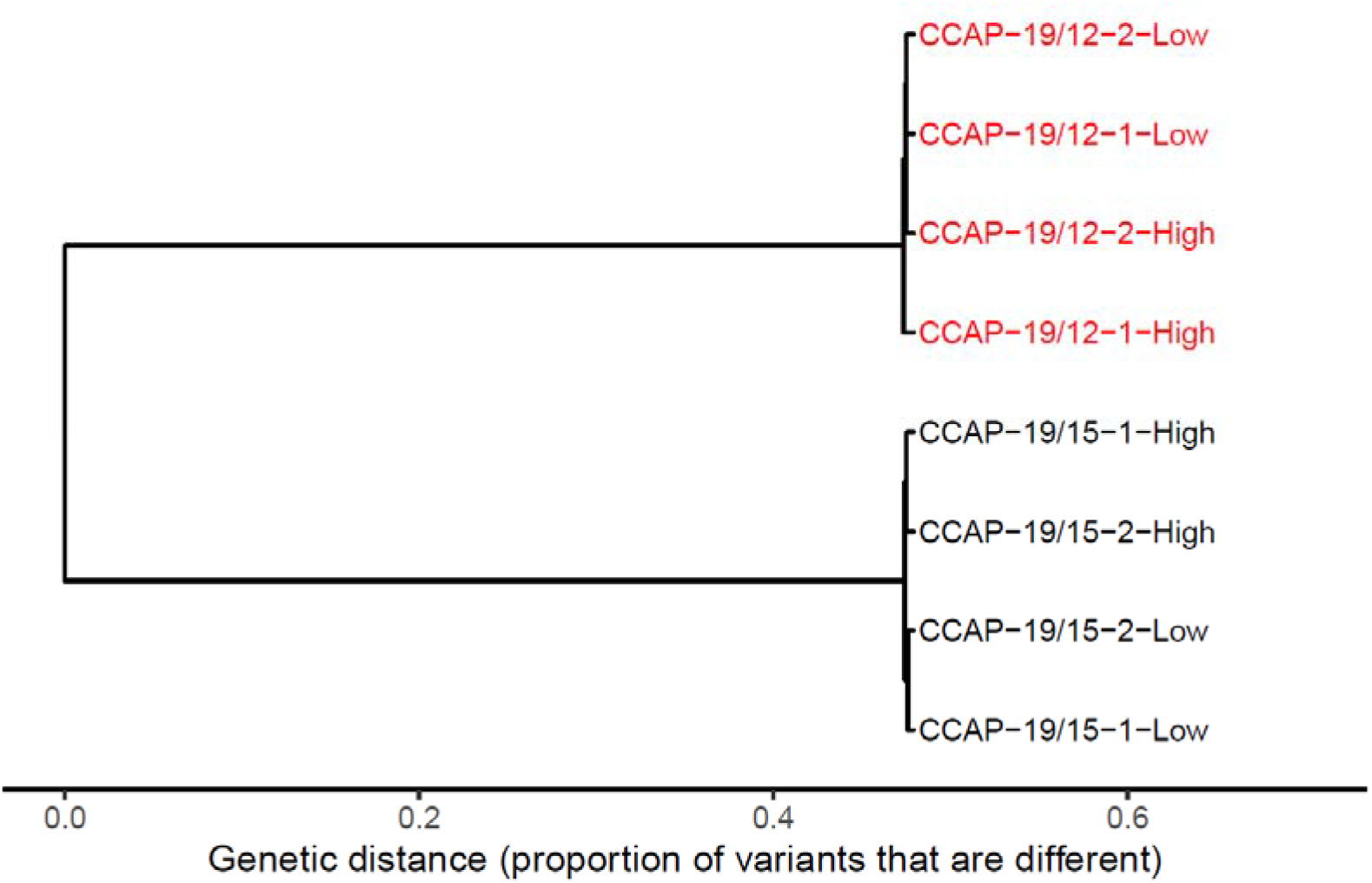
Genetic differences between the strains. Genetic distance refers to proportion of genetic differences among samples, across all 6,201 variants detected in the transcriptome. Low and High refer to salinity and numbers to samples.

### 3.2. Strains had markedly different demographic responses to osmotic stress

The two strains displayed very different population dynamics in response to osmotic stress. On the first day following transfer to a new salinity, both strains showed significantly positive growth in [NaCl] = 0.8 M and 2.4 M, but significant decline at [NaCl] = 4.0 M (Fig. 2, GLMs #1_19/12 and #1_19/15 in Table S2). However, this decline in hyperosmotic conditions was much more pronounced for CCAP 19/12 compared to CCAP 19/15, as evidenced by the highly significant positive Day:Salinity_4.0M:Strain_19/15 term (Table 1, GLM #1). While CCAP 19/12 declined by almost 80% (exp(−1.537) = 0.215 in GLM #1_19/12 in Table S2), CCAP 19/15 declined only by about 35% (exp(−0.435) = 0.647 in GLM #1_19/15 in Table S2). Strikingly, both strains then recovered from this initial decline at high salinity, and started growing again after Day 1, reaching a phase of exponential growth until *c*. Day 4 (Fig. 2). But in this phase also, the dynamics markedly differed between the strains. Notably, CCAP 19/15 grew significantly slower than CCAP 19/12 at [NaCl] = 4.0 M (highly significant negative Day:Salinity_4.0M:Strain_19/15 term in Table 1, GLM #2). In fact, the exponential growth rate of CCAP 19/12 did not significantly differ among salinities (GLM #2_19/12, Table S2), while that of CCAP 19/15 was significantly lower in [NaCl] = 2.4 M and 4.0 M as compared to 0.8 M (GLM #2_19/15, Table S2). These dynamics led to a negative relationship between initial growth rate and later exponential growth rate at high salinity, with the two strains occupying different regions along this relationship (rapid decline and growth for CCAP 19/12, slow decline and growth for CCAP 19/15), while no such pattern was observed at lower salinities (Fig. 2B). We further showed that the faster growth rate of CCAP 19/12 during the exponential phase was not explained by released density-dependent competition as a result of its drastic initial decline (from GLM #2_19/12 and GLM #2_19/15, and GLM #3, Table S2), and that the dynamics of decline and rebound was not compatible with evolutionary rescue (Gomulkiewicz & Holt 1995; Bell & Gonzalez 2009), as it also occurred in isogenic populations (Fig. S2 and Table S3).

**Fig. 2.**
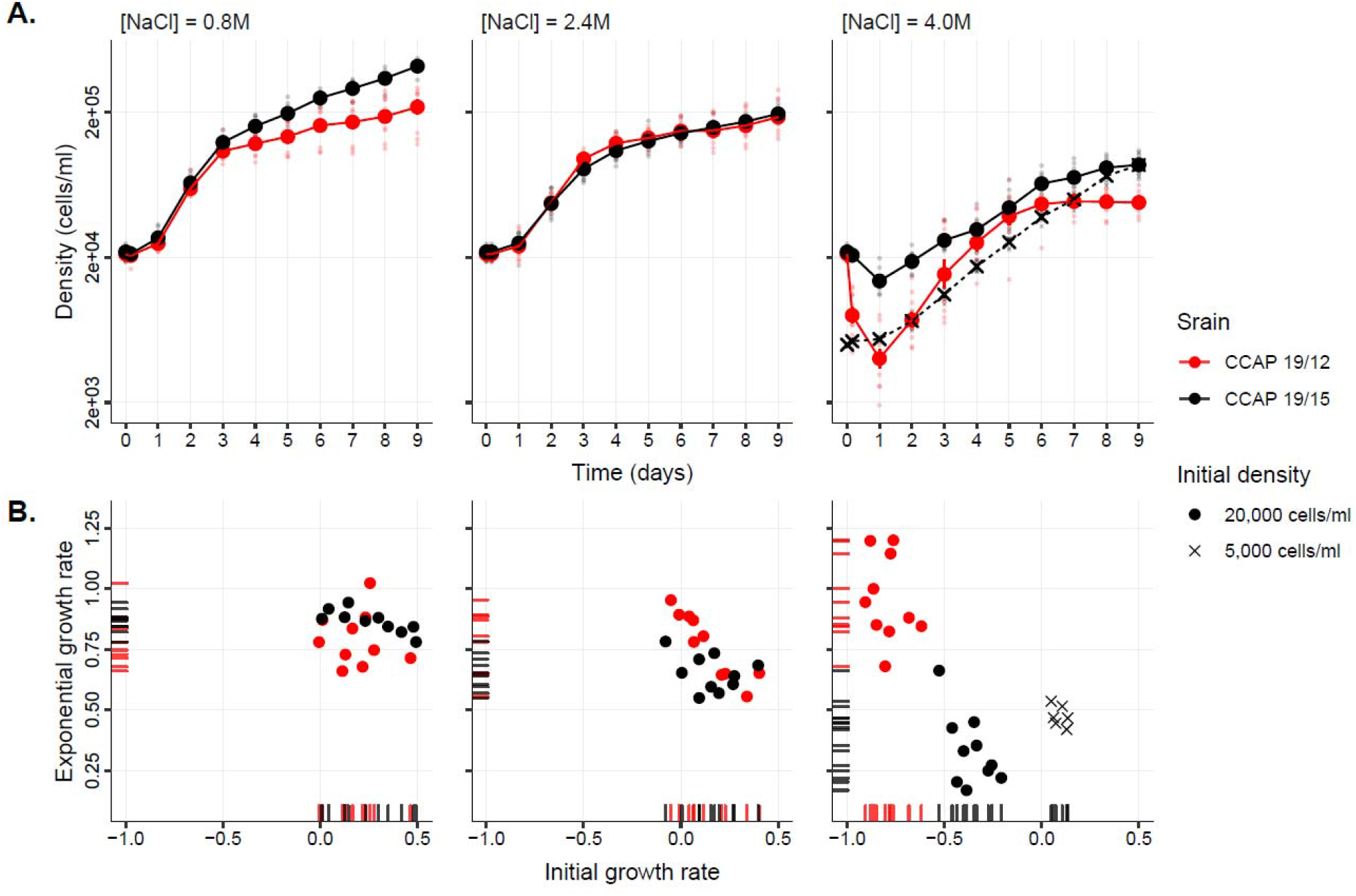
Population dynamics of two *Duniella salina* strains under different osmotic stresses. **A**. Mean population growth curve in three different salinities. For each strain (CCAP 19/12 and CCAP 19/15 in red and black, respectively), mean cell density and standard error were calculated for the experiments starting with an initial density of 20000 cells × ml^-1^ (circle) for both strains (10 distinct populations for each), or 5000 cells × ml^-1^ (cross) for CCAP 19/15 only (6 populations). **B**. Relationship between the initial and exponential growth rates following different osmotic stresses. Rug plots illustrate the distribution of the initial (days 0 to 1) and exponential (days 1 to 4) growth rates on their respective axes.

**Table 1.** Strain and salinity effects on population growth. General Linear Models (GLMs) with a negative binomial distribution were performed on cells count data, where the interaction of time (Day) with salinity or strain estimates effects of the latter on population exponential growth (or decline) rates.

|  | Estimate | Std. Error | z value | Pr(>\|z\|) |  |
| --- | --- | --- | --- | --- | --- |
| ***GLM #1: Initial population response (days 0 to 1), starting from same initial density*** |  |  |  |  |  |
| Intercept (Strain_19/12 at Salinity 0.8 M and Day 0) | 9.925 | 0.022 | 458.649 | < 0.001 | *** |
| Day | 0.197 | 0.072 | 2.723 | 0.006 | ** |
| Day:Salinity_2.4M | -0.024 | 0.096 | -0.254 | 0.800 |  |
| Day:Salinity_4.0M | -1.797 | 0.096 | -18.673 | < 0.001 | *** |
| Day:Strain_19/15 | 0.096 | 0.096 | 0.994 | 0.320 |  |
| Day:Salinity_2.4M:Strain_19/15 | -0.048 | 0.136 | -0.349 | 0.727 |  |
| Day:Salinity_4.0M:Strain_19/15 | 1.135 | 0.136 | 8.340 | < 0.001 | *** |
| ***GLM #2: Population growth in exponential phase (days 1 to 4)*** |  |  |  |  |  |
| Intercept (Strain_19/12 at Salinity 0.8 M and Day 1) | 10.317 | 0.071 | 145.898 | < 0.001 | *** |
| Day | 0.541 | 0.038 | 14.305 | < 0.001 | *** |
| Salinity_2.4M | -0.134 | 0.100 | -1.337 | 0.181 |  |
| Salinity_4.0M | -1.904 | 0.100 | -19.032 | < 0.001 | *** |
| Strain_19/15 | 0.056 | 0.100 | 0.565 | 0.572 |  |
| Day:Salinity_2.4M | 0.018 | 0.053 | 0.340 | 0.734 |  |
| Day:Salinity_4.0M | 0.095 | 0.053 | 1.784 | 0.075 |  |
| Day:Strain_19/15 | 0.056 | 0.053 | 1.043 | 0.297 |  |
| Salinity_2.4M:Strain_19/15 | -0.021 | 0.141 | -0.151 | 0.880 |  |
| Salinity_4.0M:Strain_19/15 | 1.095 | 0.141 | 7.741 | < 0.001 | *** |
| Day:Salinity_2.4M:Strain_19/15 | -0.118 | 0.076 | -1.566 | 0.117 |  |
| Day:Salinity_4.0M:Strain_19/15 | -0.411 | 0.076 | -5.436 | < 0.001 | *** |

### 3.3. Gene expression responses to osmotic stresses

Structural gene annotation resulted in 31,926 putative transcripts present across the eight samples. PCA on the total gene expression variation revealed that samples first clustered according to the strain of origin (first PC axis, 61% of variance, Fig. 3A), and then according to the salinity treatment (2^nd^ PC axis, 17% of variance, Fig. 3A). This result was confirmed by the detection of almost twice as many differentially expressed (DE) transcripts between strains (n = 3122, Fig. 3B and 3C) as between salinities (n = 1659, Fig. 3B and 3D). In addition, both strains also displayed strain-specific DE transcripts between salinities (Fig. 3B), confirmed by a significant *strain × salinity* interaction (n = 2199, Fig. 3E) on gene expression.

**Fig. 3.**
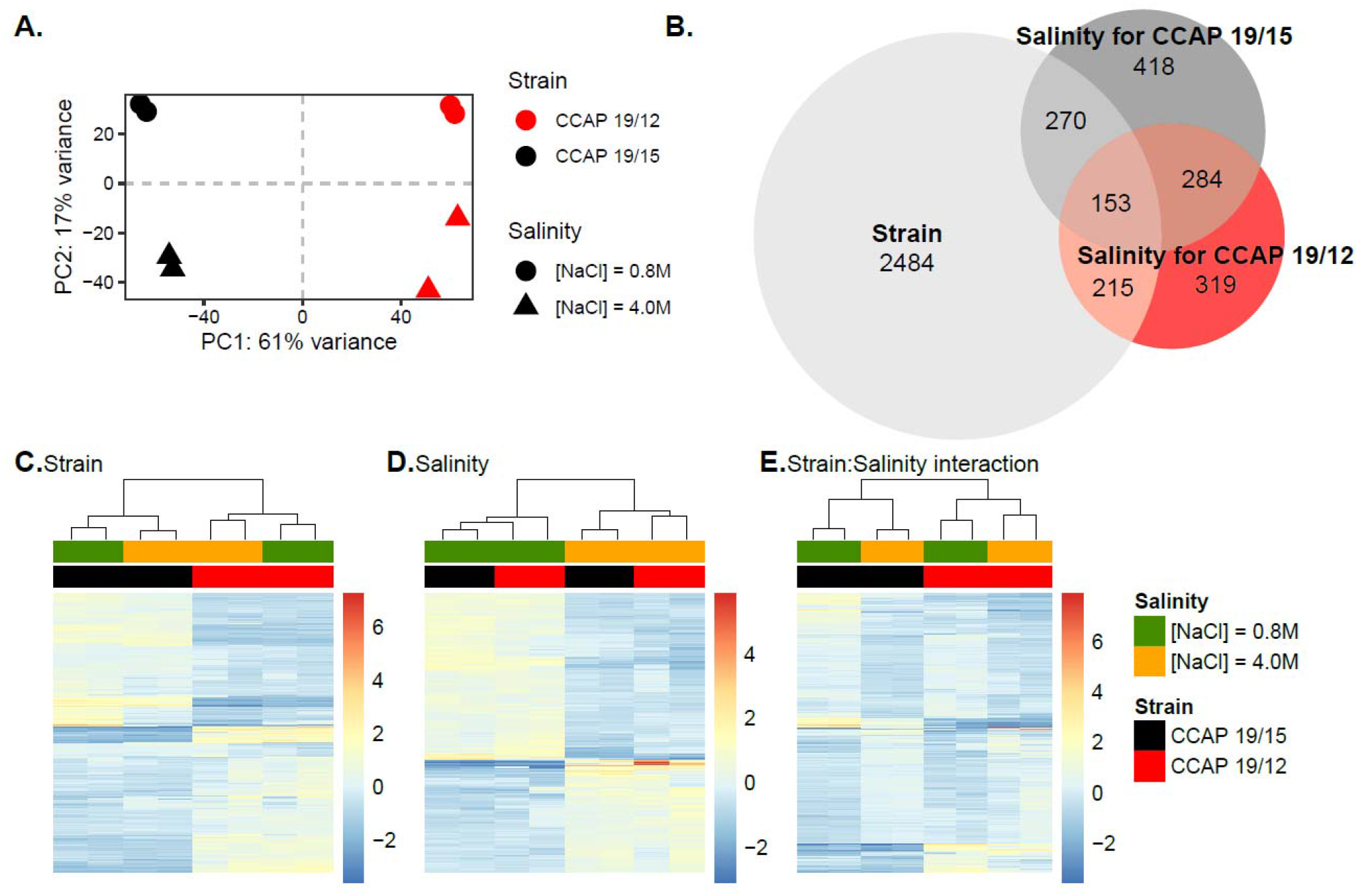
Gene expression response to osmotic stresses of two *D. salina* strains. **A**. Principal component analysis (PCA) of gene expression level for two *D. salina* strains (color) facing hypo-and hyper-osmotic stress (shape). **B**. Venn diagram showing the numbers of transcript that are significantly differentially expressed (DE; FDR < 0.05 and |Log_2_FC| > 1) between strains (light grey), salinities for CCAP 19/12 (red), or for CCAP 19/15 (dark grey). **C-E**. Heat-maps of RNA-Seq transcriptome analyses for significant DE transcripts between strains (C; n = 3122), exclusively between salinities and common to the two strains (D; n = 284), and for strain × salinity interaction, identified by performing likelihood-ratio test (LRT, FDR < 0.05) as implemented in *DESeq2* (E; n = 2199). Each row and column represent a transcript and a biological replicate, respectively. Relative expression intensities among replicates vary from blue (under-expressed) to red (over-expressed), as shown on the right-hand side of the heat-maps. Dendrograms on the top resulted from a hierarchical clustering analysis using the Euclidean distance of the relative transcript expression level among replicates.

The proportions of up- and down-regulated transcripts were similar for strain CCAP 19/12 and CCAP 19/15 (Fig. 3C). For salinity-specific DE transcripts, we observed slightly more transcripts with higher expression in low salinity than in high salinity (Fig. 3D). The strain × salinity interaction is characterized by greater gene expression differences between salinities for CCAP 19/15 than for CCAP 19/12 (Fig. 3E). For instance, we detected higher expression at low salinity for a group of transcripts, but only for CCAP 19/15 (Fig. 3E, top-left of the heat-map).

### 3.4. DNA methylation variation is structured by genomic contexts

WGB-sequencing generated a total of 4.05 × 10^8^ 150bp paired-end raw reads from eight samples. Considering *D. salina*’s genome size of *c.a*. 350 Mbp (Polle *et al*. 2017), this resulted in an estimated average depth of coverage of 43.43x (*s.d*. 3.63x) per sample (Table S1). After the data filtering, we performed our methylation analyses on an average of 5.19 × 10^8^ (*s.d*. 1.08 × 10^8^) cytosines per samples, and observed that DNA methylation is not randomly distributed along the genome (Fig. 4A). Cytosine in CpG context displayed the highest methylation level, as compared to CHG and CHH contexts. Furthermore, the nuclear genome had the most methylated cytosine, as compared to the mitochondria and chloroplast genomes (Fig. 4B). Because the highest and most variable methylation level was found at CpG methylation in our dataset, and because of their suggested role in gene regulation, while non-CpG methylation haven been described mostly as silencers for transposable elements (Law & Jacobsen 2010; Chen *et al*. 2018; Zhang *et al*. 2018; de Mendoza *et al*. 2020), we investigated the genomic DNA methylation patterns in response to salinity across the two strains only for cytosines at the CpG context.

**Fig. 4.**
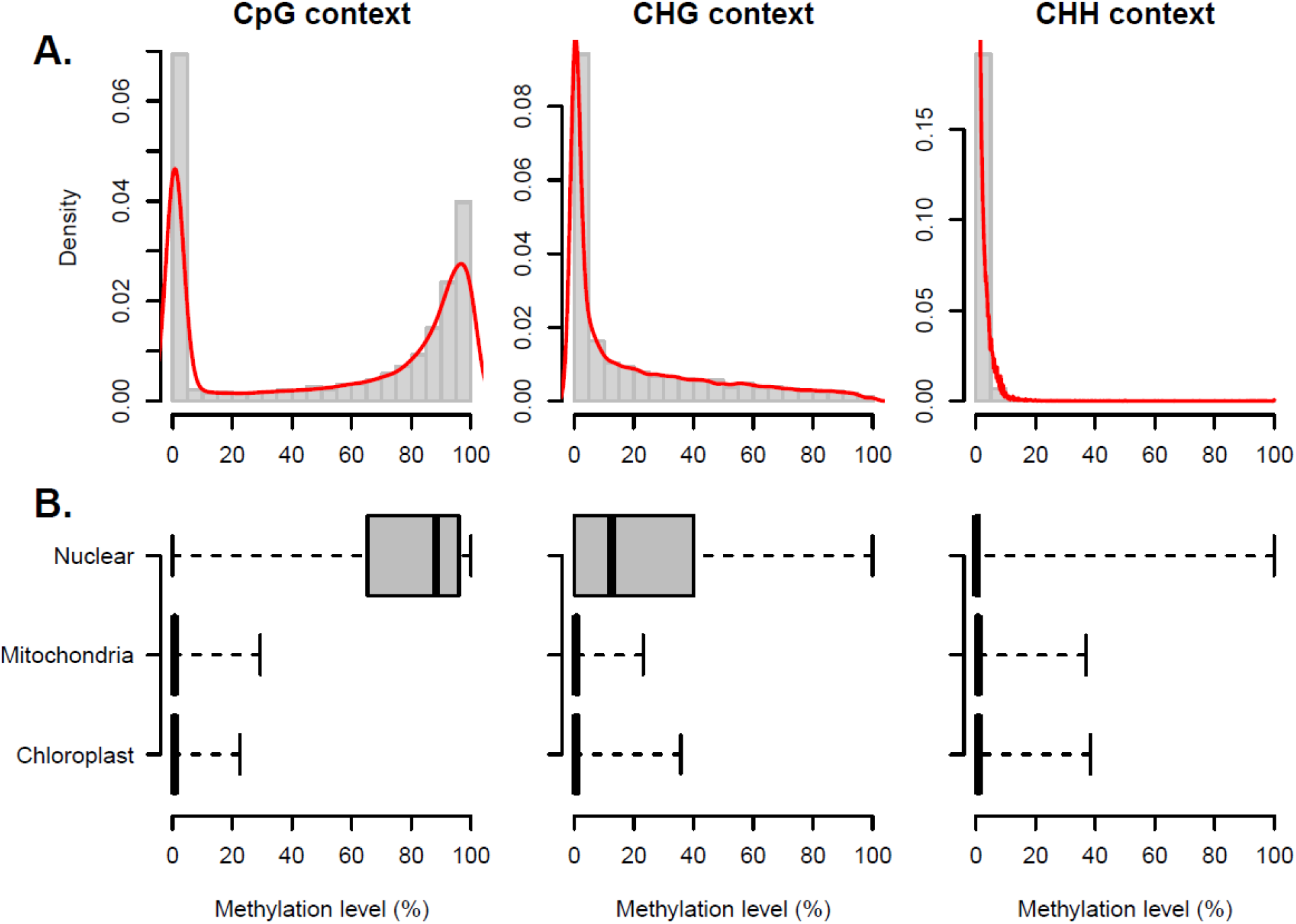
Methylation level and pattern in *Dunaliella salina*. **A**. Distribution of the methylation level according to cytosine contexts. Histogram (grey) and density (red line) plots were calculated for CpG (n = 46 800), CHG (n = 69 039) and CHH (n = 227 531) context. **B**. Distribution of methylation level for the nuclear (n = 32 409, 58 391 and 173 653 for CpG, CHG and CHH, respectively), mitochondria (n = 963, 929 and 4 381 for CpG, CHG and CHH, respectively) and chloroplast (n = 13 428, 9 719 and 49 497 for CpG, CHG and CHH, respectively) genomes, for each cytosine context.

Redundancy analyses revealed a significant effect of strain (*R*^*2*^ = 58.69%; *P* = 0.005) on total DNA methylation at the CpG context. However, contrary to gene expression, we did not detect any significant marginal effect of salinity (*R*^*2*^ = 6.47%; *P* = 0.602) on epigenetic state, as also observed on the PCA plot (Fig. 5A). On a coarser scale, considering non-overlapping 100bp windows, we detected 932 DMRs between the two strains, but only 27 DMR between salinities when all strains were pooled together, and no significant strain × salinity interaction (*R*^*2*^ = 5.83%; *P* = 0.566). Nevertheless, we did detect a few strain-specific DMRs between salinities for each strain (n = 40 and 54, for CCAP 19/12 and 19/15 respectively; Fig. 5B to 5E).

**Fig. 5.**
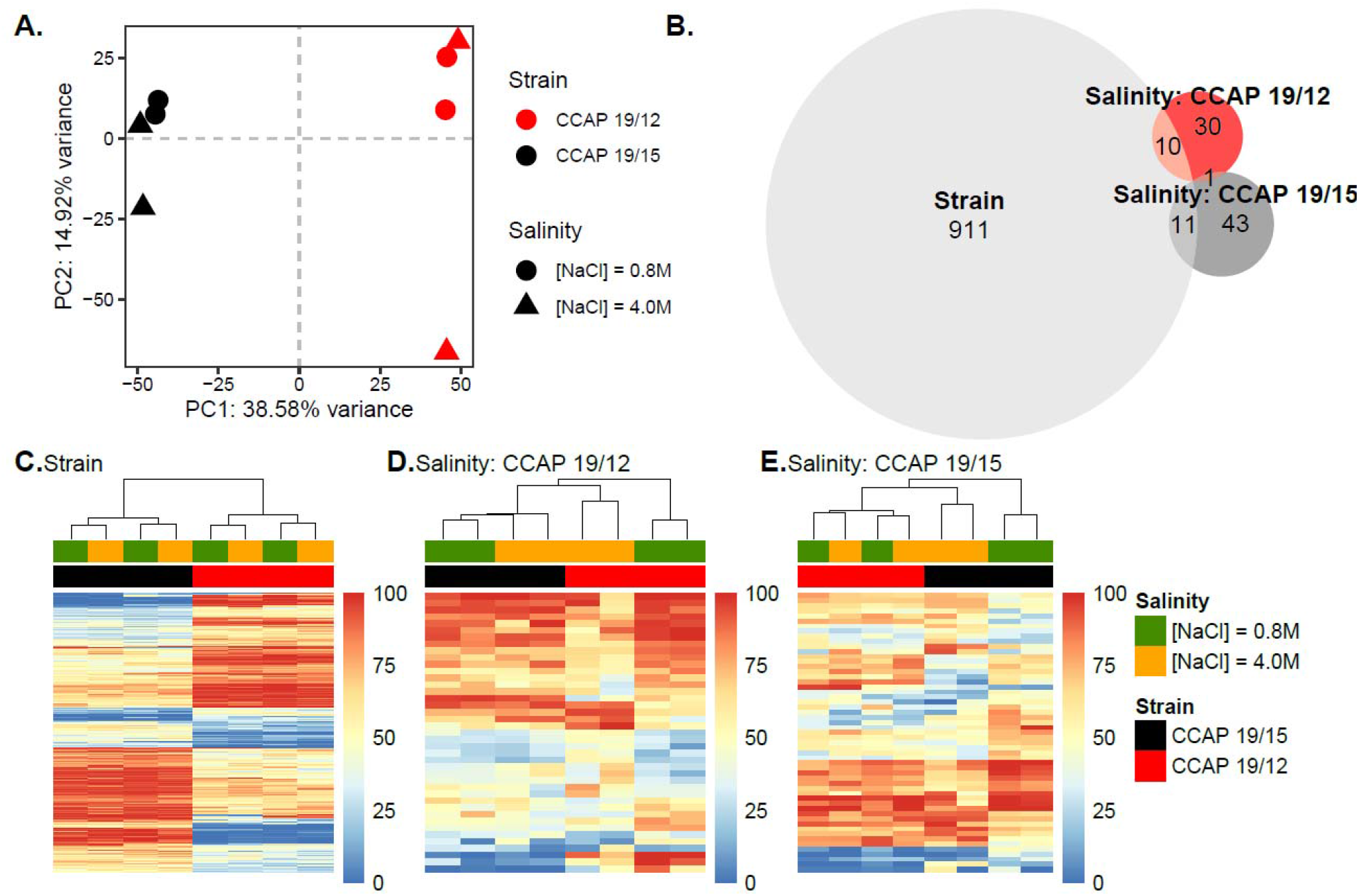
DNA methylation response to osmotic stresses of two *D. salina* strains. **A**. Principal component analysis (PCA) of DNA methylation level for two *D. salina* strains (color) facing hypo- and hyper-osmotic stress (shape). **B**. Venn diagram showing the numbers of differentially methylated regions (DMR; *q*-value < 0.05 and |diff-Methylation| > 20%) between strains (light grey), salinities for CCAP 19/12 (red), or for CCAP 19/15 (dark grey). **C-E**. Heat-maps of WGB-Seq analysis for DMRs between strains (C; n = 911), or between salinities for CCAP 19/12 (D; n = 40) and CCAP 19/15 (E; n = 54). Each row and column represent a DMRs and a biological replicate, respectively. DNA methylation levels vary from blue (unmethylated) to red (methylated), as shown on the right-hand side of the heat-maps. Dendrograms on the top resulted from a hierarchical clustering analysis using the Euclidean distance of DNA methylation level among replicates.

### 3.5. Gene expression depends on both the genotype and the environment

The genotype, the environment, and the CpG methylation level jointly significantly explained an important part of total transcriptomic variation (adjusted *R*^*2*^ = 88.11%; *P* = 0.005; Fig. 6A). More precisely, the strong correlation between the genotype and DNA methylation reported above (*R*^*2*^= 58.69%; *P* = 0.005) resulted in a great confounding effect (*R*^*2*^ = 62.73%), whereby the influence of genotype and epigenetic state on gene expression could not be disentangled. Nevertheless, we still detected significant marginal effects of both the environment (adjusted *R*^*2*^ = 17.43%; *P* = 0.022) and CpG methylation (adjusted *R*^*2*^ = 9.51%; *P* = 0.043) on transcriptomic variation (Fig. 6A). These results indicated a larger role of the genotype on gene expression, but also underlined the epigenetic processes and the environment as additional sources of variation in gene expression.

**Fig. 6.**
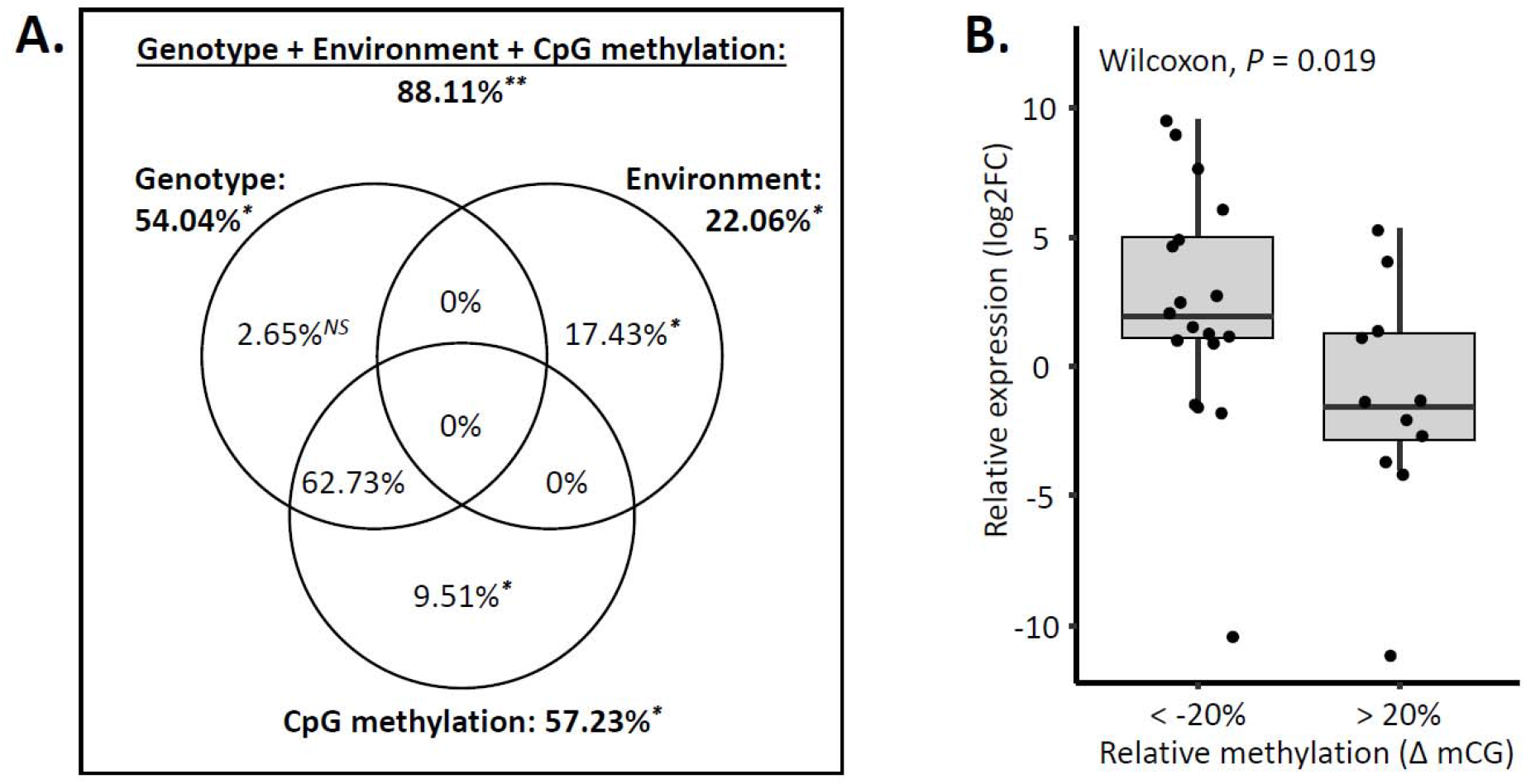
Correlation between gene expression and epigenetic variation. **A**. Transcriptomic variation partitioning was performed based on Borcard *et al*. (1992). Proportions of the total transcriptomic variation explained by the genotype (strains CCAP 19/12 vs CCAP 19/15), the environment (low vs high salinities, controlling for paired samples) and cytosine methylation on CpG context are based on the adjusted *R*^*2*^ of RDA analyses. Bold number outside circles refer to total percentage of variation (adjusted *R*^*2*^) explained by the genotype, the environment and CpG methylation (top, full model), or by only the genotype, the environment or CpG methylation (reduced models). Percentages inside circles indicate pure contributions to gene expression level resulting from partial RDAs that isolate the effect of each single explanatory variable, but taking into account others as covariates. Percentages within intersections indicate shared contribution across different variables to gene expression variation. ANOVA-like permutation tests were calculated on total and pure contributions. *NS*: Non-significant; ^*^: *P* < 0.05; ^**^: *P* < 0.01. **B**. Differential transcript expression against differential DNA methylation. Boxplot showing the level of expression differentiation (Log2 Fold Change) against the difference in methylation level (ΔmCG). Boxplot and mean comparison analysis were performed for only DMRs detected as significantly differentially methylated between strains or salinities, and associated to a given transcript that also showed significant differential expression between strains or salinities. Differences between strain (n = 27), between salinities for CCAP 19/12 (n = 1), between salinities for CCAP 19/15 (n = 2).

Finally, we assessed whether DNA methylation in *cis* could influence gene expression. By grouping cytosines according to the nearest TSS, we obtained 3,064 methylated regions, among which we detected 250 DMRs: 223 DMRs between the two strains, and 10 and 21 DMRs between salinities for CCAP 19/12 and 19/15, respectively, with 4 common DMRs between strain and salinity comparisons. We observed the same proportion of DE transcripts in the methylated and differentially methylated regions (χ^*2*^ = 0.420, df =1, *P* = 0.517): among the 3,058 methylated regions, 419 were associated to DE transcripts, while among the 250 DMRs, 30 were associated to significant DE transcripts for the same comparisons (i.e. between strains, or between salinities for both strain). Comparison of the relative expression level against the relative DNA methylation level revealed that hypomethylated DMRs (ΔmCG < -20%) were associated to overexpressed transcripts (Log2FC > 0), and conversely, hypermethylated (ΔmCG > 20%) DMRS showed a lower transcript expression level (Fig. 6B).

## 4. Discussion

We investigated how genetic variation relates to plastic responses at multiple biological levels, by comparing DNA methylation patterns, gene expression levels, and demographic phenotypes across salinities, in two strains of a halotolerant eukaryote known to be predominantly under selection for salinity tolerance in its natural habitat (Kirst 1990; Oren 2005; Ben-Amotz *et al*. 2009). Our results shed light on the broad molecular mechanisms of phenotypic plasticity and its genetic variation.

### 4.1. Variation of reaction norms between genotypes

We first showed that the two strains used in our study displayed distinct phenotypic response to environmental change, both in term of population dynamics and gene expression level. We observed strain-specific growth rates within the first 24h following an osmotic stress, and later during exponential growth. Specifically, strain CCAP 19/12 displayed a greater initial population decline as compared to CCAP 19/15, when cells were subjected to hyper-osmotic stress. This is possibly caused by a variable intensity of programmed cell death, a phenomenon that is widely present in unicellular microalgae, and can be triggered by a variety environmental stresses, including an increase in salinity (Zuppini *et al*. 2010; Bidle 2016; Durand & Ramsey 2019). Interestingly, the exponential growth rate following initial decline was also strain-specific, rather than only resulting from variable competition levels following different levels of decline among strains. Indeed at high salinity, CCAP 19/12 experienced faster growth than CCAP 19/15, even when the latter strain started at low density.

At the cellular level, the response of *Dunaliella salina* to osmotic stress is characterized by immediate changes in cell volume and intracellular ions, followed by slower changes in gene expression, starting within 4h to 24h under the osmotic stress (Chen & Jiang 2009), and regulating notably glycerol metabolism (Zhao *et al*. 2013). In this study, we detected differentially expressed transcripts between salinities 24h after the osmotic stress, confirming *D. salina* gene-expression plasticity in response to salinity. As we measured a very low within-strain genetic variation, we are confident that observed phenotypic differences among salinities, both in term of population dynamics and gene expression, resulted from phenotypic plasticity, rather than from selection in a heterogeneous cell population. This conclusion was also confirmed by the observation of the same population dynamics in isogenic populations.

Interestingly, we showed that responses to osmotic stress partly involved the regulation of strain-specific genes, which thus likely underlies the observed strain-specific population dynamics. We were able to successfully assign only *c*. 30% of the total transcripts to at least one gene ontology term (Fig. S3). This transcript annotation analysis revealed that differentially expressed genes are mostly related to chloroplast structure and activities (Fig. S3), but located in the nuclear genome. Interestingly, this lends support to the hypothesis above that the rapid decline followed by rebound that we observed under hyper-osmotic stress may involve chloroplast-mediated programmed cell death (PCD), as studies have underlined the potential roles of cytochrome f, responsible for oxygenic photosynthesis, and thylakoid membrane complexes, in PCD for both plants and unicellular organisms (Zuppini *et al*. 2009; Thamatrakoln *et al*. 2013; Murik *et al*. 2014; Ambastha *et al*. 2015; Bidle 2016).

Our aim was to better understand how a given phenotype is connected to different levels of the genotype-phenotype (GP) map, by empirically relating gene expression to the expression of a higher phenotype. However, explanatory power in the GP map is necessarily limited by the level with the lowest dimensionality, and here we only analyzed a few higher-level phenotypes chosen for their ecological meaningfulness. Achieving satisfying resolution in the GP map would probably require characterizing the phenotype more finely, as well as using more environmental contrasts (Chevin *et al*. 2021). For example, quantifying different chloroplast-related phenotypes (e.g. reactive oxygen species production, photosystem activity, etc.) over a gradient of salinity would help break down correlations in the plastic responses, as well as elucidating the sequence of molecular events associated with chloroplasts influencing the rate of cellular death.

### 4.2. DNA methylation patterns and gene silencing in Dunaliella salina

The levels and patterns of DNA methylation are known to vary drastically among organisms, including within microalgae (Feng *et al*. 2010; Zemach *et al*. 2010), such that characterizing these epigenetic traits and their potential roles in gene regulation remains an important goal, especially for organisms with special ecological or biological interest. Here, we present the first study of whole-genome DNA methylation in *D. salina*, a model organism for salinity tolerance, and a biotechnologically important species for carotene production (Ben-Amotz *et al*. 2009). We found that cytosine methylation is nearly absent in both the chloroplast and the mitochondrial genomes, and depends on the genomic context in the nuclear genome. More specifically, we observed the highest methylation levels of nuclear cytosines in the CpG context, but with a highly bimodal distribution, including many low methylated sites. We detected a lower degree of methylation (*c*. 10%) in the CHG context, and quasi-absence of methylation in the CHH context. Interestingly, this pattern is very similar to that observed in *Arabidopsis thaliana* (Cokus *et al*. 2008; Feng *et al*. 2010; Zhang *et al*. 2020), but quite different from those in more close related green algae such as *Chlamydomonas reinhardi* or *Volvox carteri*, which display very low methylation levels, or *Chlorella variabilis*, where genes are universally methylated (Feng *et al*. 2010; Zemach *et al*. 2010). This confirms that the type and extent of DNA methylation varies at relatively low phylogenetic scales (Bewick *et al*. 2017; Alonso *et al*. 2019).

Beyond levels and patterns, the roles of DNA cytosine methylation in microalgae remains poorly understood, and could be very variable. For example, *C. variabilis* methylation level in the CpG context in promoters was detected to be inversely correlated to gene expression, suggesting that promoter-proximal methylation represses transcription (Zemach *et al*. 2010). At the opposite, only a weak negative correlation between promoter methylation and gene transcription was observed in *V. carteri*, while CpG methylation is enriched in transposons (Zemach *et al*. 2010). In *C. reinhardtii*, cytosine methylation, observed in the chloroplast genome during gametogenesis, was suggested to be involved in the uniparental inheritance of mating type (Sager & Grabowy 1983; Umen & Goodenough 2001; Nishiyama *et al*. 2002; Nishiyama *et al*. 2004). Here, we could associate only few differentially methylated regions with differentially expressed transcripts. Nonetheless, we showed a negative correlation between CpG methylation level for TSS-proximal cytosine and gene expression, suggesting that DNA methylation at the promoters represses transcription. Methylation at CHG context remained however unclear, but could be involved in the silencing of repetitive sequences and transposon, as in land plants or *C. variabilis* (Saze & Kakutani 2011; Kim *et al*. 2015b).

### 4.3. Sources of phenotypic variation and plasticity

One of the aims of our study was to quantify the relative importance of genetic and non-genetic (environmental and epigenetic) sources of gene expression. We detected a substantial contribution of the genotype to both the epigenetic and phenotypic variation. As we also observed a negative correlation of DMRs and DE transcript, these results suggest that DNA methylation at CpG context is an intermediate step between the genotype and the gene-expression phenotype. However, the absence of a significant marginal effect of salinity or strain × salinity interaction on epigenetic state questioned the role of individual CpG methylations in plasticity. In plants and vertebrates, cytosine methylation responsible for gene expression regulation is mostly organised in specific regions, such CpG islands, frequent in the promoters of transcript start sites, or within coding sequences (Bird 2002; Jaenisch & Bird 2003; Zilberman *et al*. 2007; Law & Jacobsen 2010; Deaton & Bird 2011; Gallusci *et al*. 2016), while the functional relevance of single CG methylations remain ambiguous (Denkena *et al*. 2021). Here, we did detected strain-specific DMRs between salinities, but only when considering 100bp non-overlapping regions, rather than individual CpG sites. This result underlined that methylation of CpG regions (rather than individual sites) are involved in gene expression plasticity and its genetic variation. Furthermore, the strain-specific epigenetic responses to environmental conditions highlight the evolutionary potential of epigenetic plasticity.

We found that DNA methylation levels vary little across salinity, and also co-vary little with gene expression is *cis*. For this to be consistent with an important role of DNA methylation in observed levels of gene expression across salinity, we must hypothesise that methylation-mediated *cis*-regulation at a small number of genes is enough to cause *trans*-regulation of expression of many other genes. This hypothesis is supported by the significant marginal environmental effect that we detected on gene expression levels, and by the negative correlation between DNA methylation and gene expression for a few genes. Alternatively, the responses to osmotic stress that we observed may be too rapid to be explained by DNA methylation. DNA methyltransferases establish DNA methylations during cell divisions (Law & Jacobsen 2010), which occur roughly once per day in *D. salina* (Ben-Amotz *et al*. 2009), so assessing DNA methylation 24h after an osmotic stress as we did here should in theory be sufficient to observe epigenetic responses to salinity. However, the cell cycle is likely to depend on salinity and the strain, as reflected by their demographic dynamics (Figure 2). Other mechanisms of gene expression regulation are widespread in microalgae, such as posttranslational histone modifications, and small-RNA mediated pathways (Kim *et al*. 2015b), which could also be involved the gene expression plasticity we observed in *D. salina*.

Although we detected that gene expression variation is mostly explained by the genotype and CpG methylation, we could not disentangle their influences, due to their strong correlation. Nevertheless, the significant genotype × environment interaction on gene expression highlights the evolutionary potential of plasticity, as the environment affects distinct genotypes differently, and in a multi-level manner.

In conclusion, we confirmed that the molecular mechanisms contributing to phenotypic plasticity in the halotolerant microalga *Dunaliella salina* involve variation in DNA methylation and gene expression with the environment, but found relatively weak contribution of DNA methylation to gene expression. Importantly, the large contribution of the genotype to the observed variation at multiple levels, including to demographic traits that are direct components of absolute fitness (Fig. 2), highlights the evolutionary potential of phenotypic plasticity at multiple molecular levels. An interesting avenue for future research will be to analyze the experimental evolution of plasticity at different levels, from DNA methylation to gene expression to higher phenotypes.

## Acknowledgements

This work was supported by the European Research Council (Grant 678140-FluctEvol) to LMC, and a Fonds de Recherche du Québec - Nature et Technologies (FRQNT) fellowship to CL.

## Statement of authorship

CL conceived and designed the study, with inputs from DG and L-MC. CL and DG performed the experiment and collected the data. CL analysed the data, prepared figures and tables, and wrote the original draft, L-MC reviewed and edited the draft. All the authors reviewed and approved the final draft of the manuscript.

## Data accessibility and Benefit-Sharing

Data accessibility: Raw sequence data (RNA-seq and WGB-seq) used in this study are deposited in the NCBI’s Sequence Read Archive (SRA) database under BioProject ID PRJNA736997. The specific samples used in this study are under the BioSample accessions SAMN19677492, SAMN19677495, SAMN19677496, SAMN19677497, SAMN19677507, SAMN19677510,

SAMN19677511 and SAMN19677512. The *de novo* transcriptome assembly and growth rate data will be deposited in a public digital repository once a decision will be made on the manuscript.

Benefits Generated: Benefits from this research accrue from the sharing of our data and results on public databases as described above.

## Supplementary Information

**Table S1.**
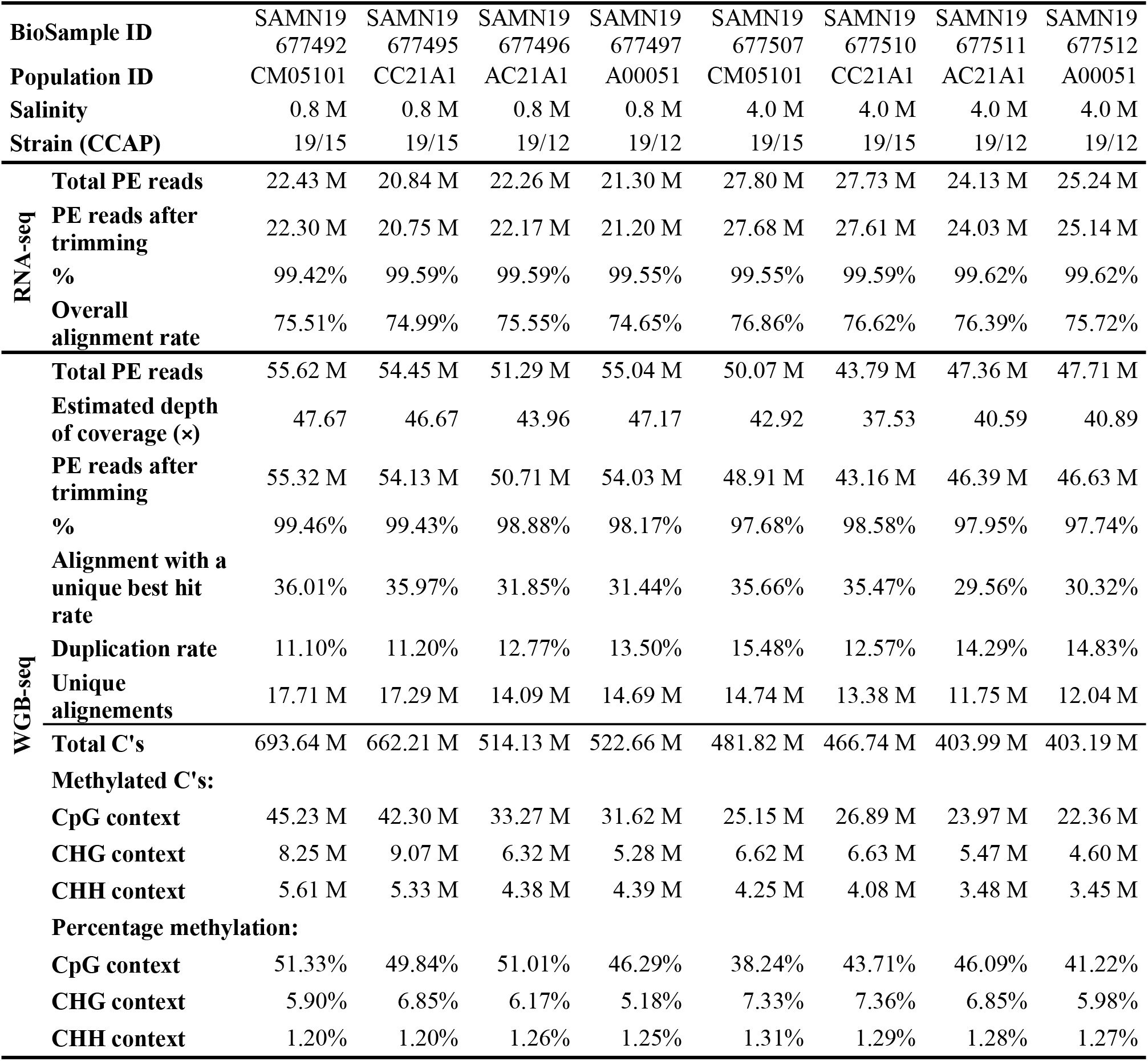
Mapping statistics for RNA-seq and WGB-seq data.

**Table S2.**
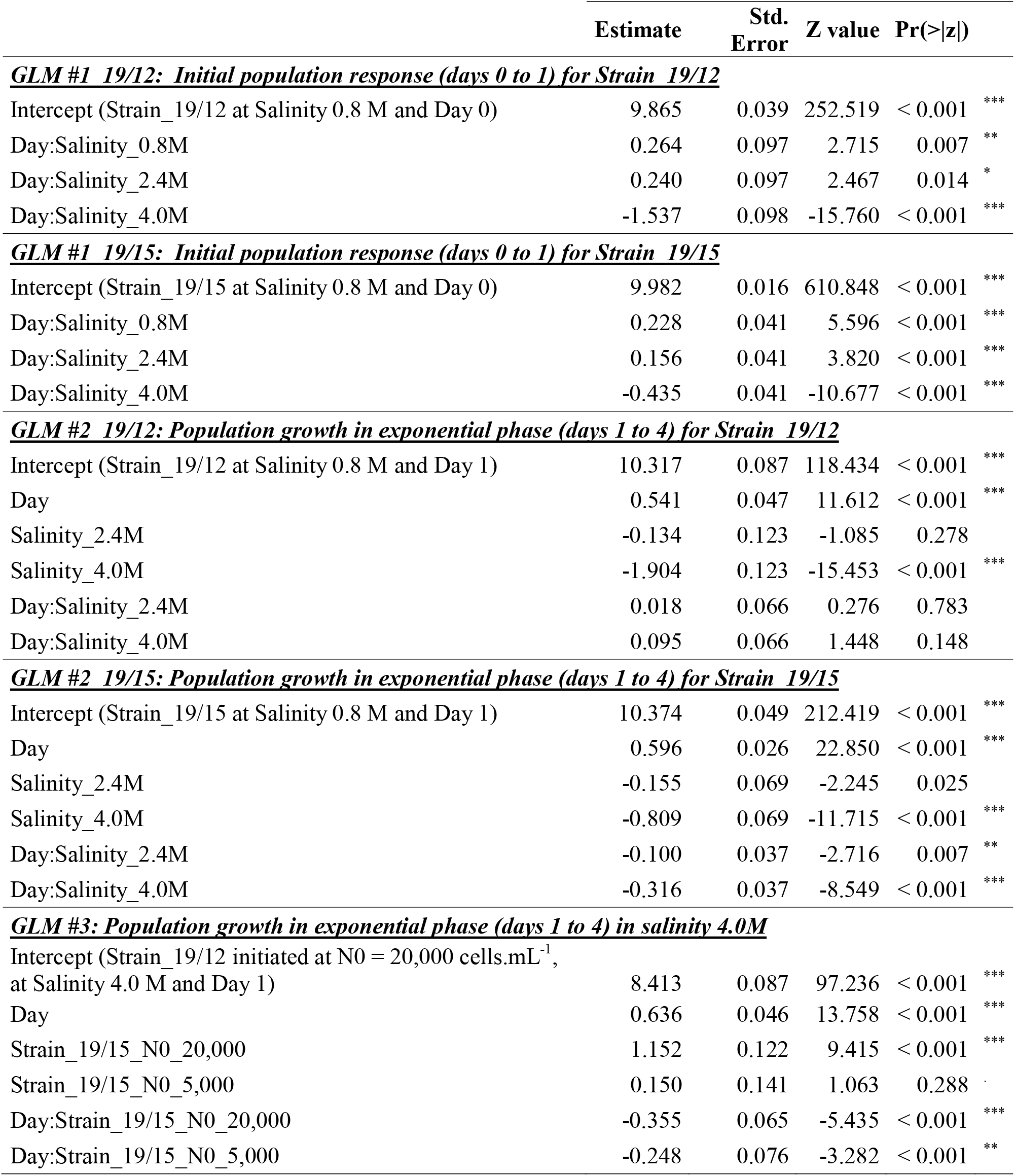
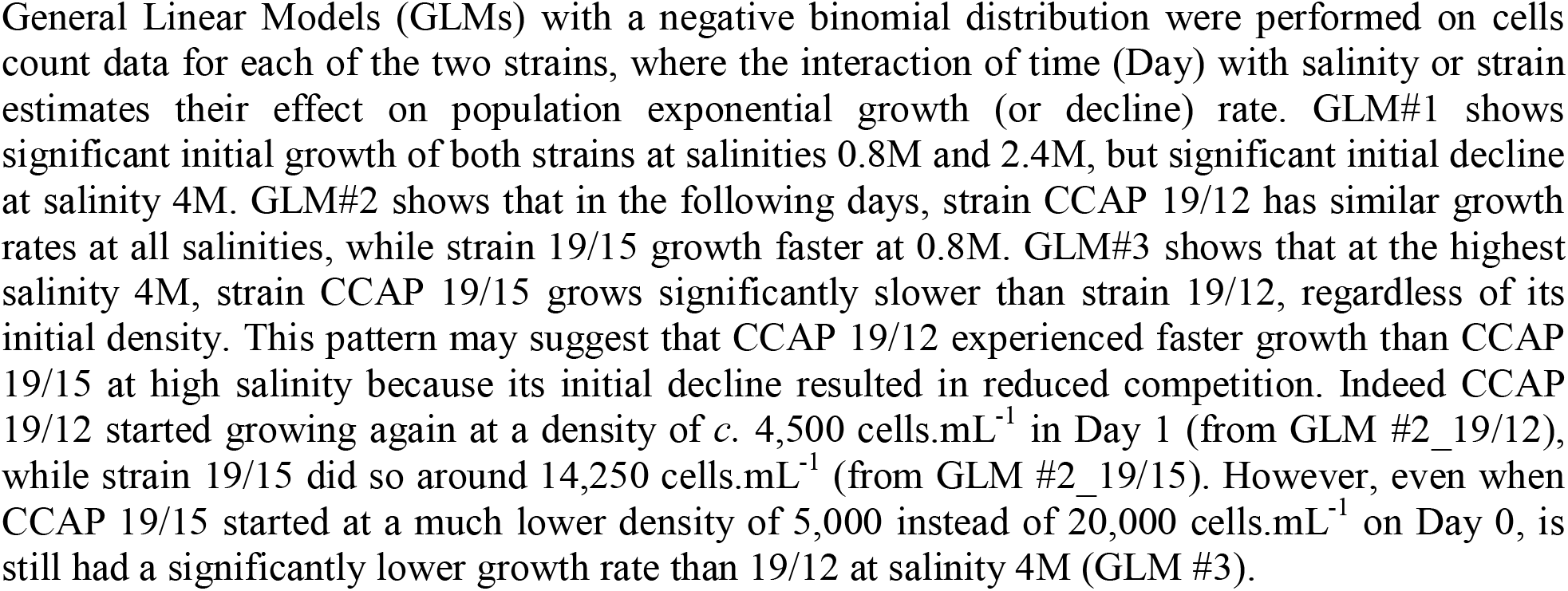
Effects of salinity or initial density on population growth rate for CCAP 19/12 and 19/15.

**Table S3.** Strain and salinity effects on population growth rate for isogenic populations.

|  | Estimate | Std. Error | z value | Pr(>\|z\|) |  |
| --- | --- | --- | --- | --- | --- |
| ***GLM #1 Isogenic: Initial population response (days 0 to 1), starting from same initial density*** |  |  |  |  |  |
| Intercept (Strain_19/12 at Salinity 0.8 M and Day 0) | 9.494 | 0.056 | 169.002 | < 2e-16 | *** |
| Day | 0.813 | 0.173 | 4.697 | 0.000 | *** |
| Day:Salinity_2.4M | -0.403 | 0.279 | -1.443 | 0.149 |  |
| Day:Salinity_4.0M | -3.586 | 0.228 | -15.696 | < 2e-16 | *** |
| Day:Strain_19/15 | -0.198 | 0.228 | -0.870 | 0.384 |  |
| Day:Salinity_2.4M:Strain_19/15 | 0.058 | 0.395 | 0.147 | 0.883 |  |
| Day:Salinity_4.0M:Strain_19/15 | 3.045 | 0.323 | 9.438 | < 2e-16 | *** |
| ***GLM #2 Isogenic: Population growth in exponential phase (days 1 to 4)*** |  |  |  |  |  |
| Intercept (Strain_19/12 at Salinity 0.8 M and Day 1) | 10.379 | 0.108 | 96.317 | < 2e-16 | *** |
| Day | 0.577 | 0.058 | 10.024 | < 2e-16 | *** |
| Salinity_2.4M | -0.411 | 0.187 | -2.202 | 0.028 | * |
| Salinity_4.0M | -3.252 | 0.153 | -21.288 | < 2e-16 | *** |
| Strain_19/15 | -0.125 | 0.152 | -0.820 | 0.412 |  |
| Day:Salinity_2.4M | 0.028 | 0.100 | 0.279 | 0.780 |  |
| Day:Salinity_4.0M | 0.362 | 0.082 | 4.434 | 0.000 | *** |
| Day:Strain_19/15 | 0.127 | 0.081 | 1.563 | 0.118 |  |
| Salinity_2.4M:Strain_19/15 | -0.091 | 0.264 | -0.343 | 0.731 |  |
| Salinity_4.0M:Strain_19/15 | 2.550 | 0.216 | 11.818 | < 2e-16 | *** |
| Day:Salinity_2.4M:Strain_19/15 | -0.080 | 0.141 | -0.570 | 0.569 |  |
| Day:Salinity_4.0M:Strain_19/15 | -0.798 | 0.115 | -6.919 | 0.000 | *** |

**Fig. S1.**
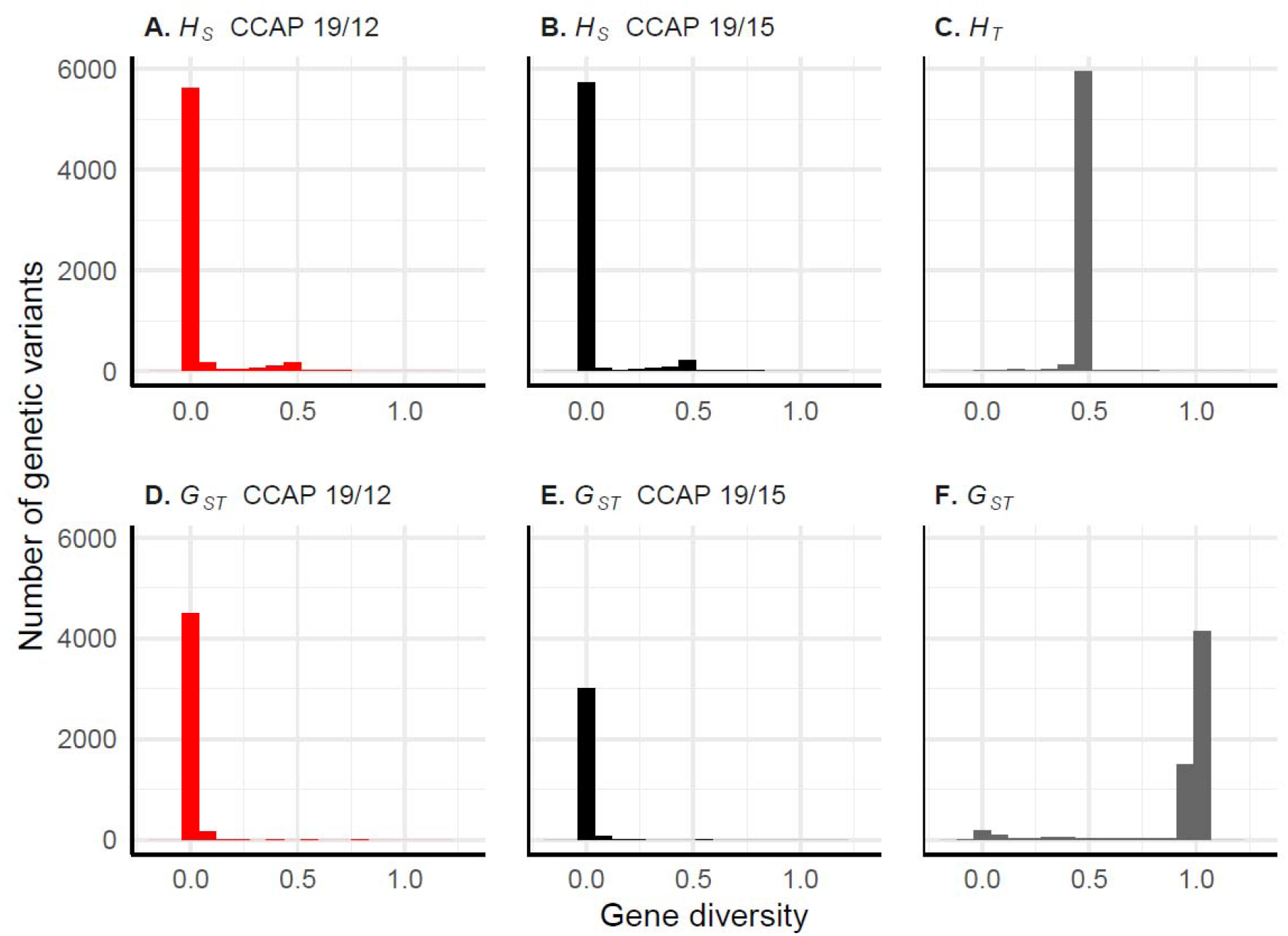
Genetic variation within and among populations. Gene diversity within population (*H*_*S*_), averaged across populations of CCAP 19/12 (A) or 19/15 (B), total gene diversity across all populations (*H*_*T*_, C), and genetic differentiation (*G*_*ST*_) across populations of CCAP 19/12 (D), of 19/15 (E), or between strains (F).

**Fig. S2.**
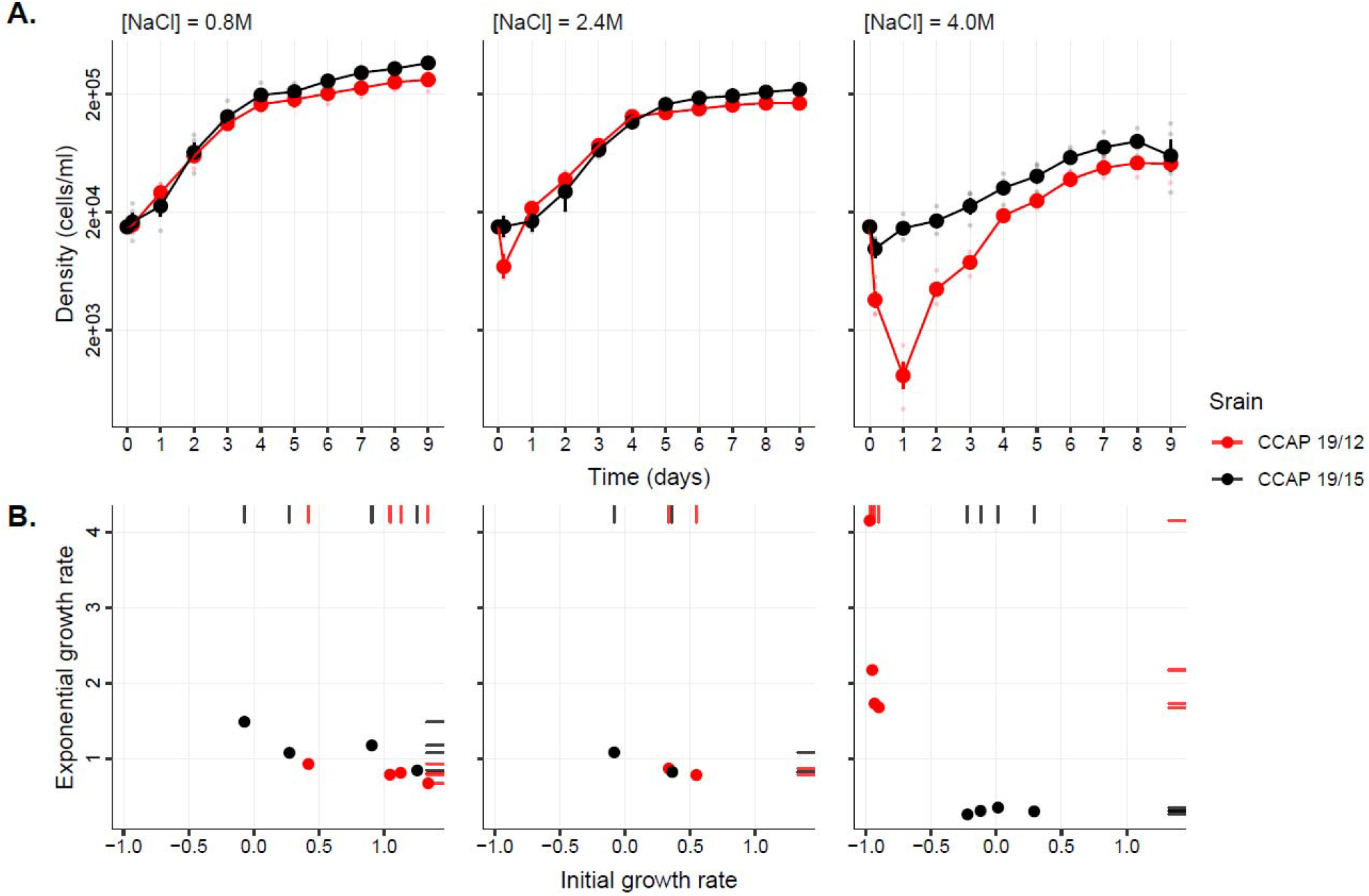
Dynamics of isogenic populations under different osmotic stresses. Isogenic populations were founded from single isolated cells for each of the two strains following the protocol in Leung *et al*. (2020). As *D. salina* is haploid, a population founded from a single cell is expected to be isogenic. **A**. Mean population growth curves in three different salinities. For each strain (CCAP 19/12 and CCAP 19/15 in red and black, respectively), mean cell density and standard error were calculated from four isogenic populations (or two for the iso-osmotic condition). **B**. Exponential growth rate against initial growth or decline rate, under different osmotic regimes. Rug plots illustrate the distribution of the initial (days 0 to 1) and exponential (days 1 to 4) growth rates on their respective axes.

**Fig. S3.**
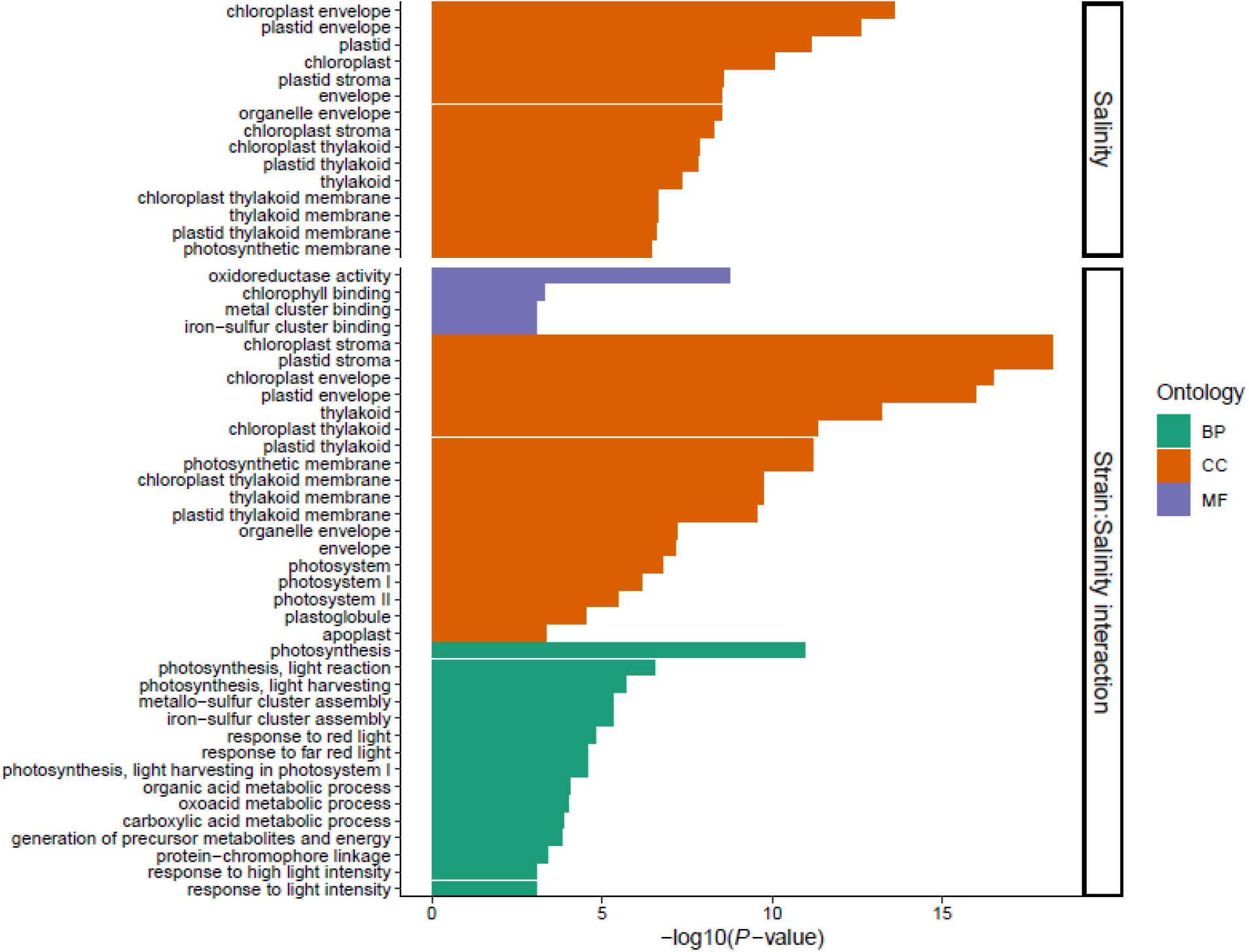
Gene ontology (GO) term enrichment analysis. We adopted GO assignments to classify the functions of *D. salina* transcripts. Based on sequence homology to known sequence databases (BLAST+/SwissProt), a total of 9,874 *D. salina* transcripts (30.93% of 31,926 transcripts) were assigned at least one GO term and categorized into 12,543 GO terms. Enriched GO terms of the DE genes were then identified using Fisher’s exact test in the *topGO* R package (Alexa and Rahnenfuhrer, 2010). GO categories included molecular function (MF, purple), cellular component (CC, orange) and biological process (BP, green),) and were sorted by decreasing order of evidence within each category, based on the GO enrichment test *P*-value (for *P* ≤ 0.001) after Benjamini-Hochberg (BH) adjustment. No significant GO term enrichment was detected for DE transcripts between strains.

